# KLHL40 gene replacement therapy in severe nemaline myopathy improves survival and skeletal muscle function in a preclinical mouse model

**DOI:** 10.64898/2026.09.23.753846

**Authors:** Grace R Tate, Joanna McNulty, Jeffrey Widrick, Vandana A Gupta

**Author notes:** To whom correspondence should be addressed: Vandana A Gupta, Division of Genetics, Department of Medicine, Brigham and Women’s Hospital, Harvard Medical School, Boston, MA 02115, USA.

## Abstract

Nemaline myopathy is a severe inherited skeletal muscle disorder characterized by profound muscle weakness, impaired mobility, and early mortality in severe cases. Despite the devastating clinical burden, no disease-modifying therapies are currently available. Nemaline myopathy-causing genes either encode sarcomeric structural proteins or regulators of sarcomeric protein stability, resulting in primary defects in sarcomere structure and function. Recessive mutations in *KLHL40* cause one of the most severe forms of nemaline myopathy, frequently resulting in perinatal lethality. Given its loss-of-function mechanism, KLHL40-associated nemaline myopathy is an attractive candidate for gene replacement therapy. We therefore evaluated the therapeutic potential of systemic AAV9-mediated *KLHL40* gene replacement in a mouse model of severe nemaline myopathy. A single systemic administration of an AAV9 vector encoding human KLHL40 during early disease produced durable rescue of survival, skeletal muscle pathology, motor performance, and contractile function in KLHL40-deficient mice. Therapeutic benefit was accompanied by restoration of sarcomere organization and normalization of myofiber size without detectable long-term histopathological abnormalities in major organs. Collectively, our findings provide a strong preclinical foundation for developing AAV9-mediated *KLHL40* gene replacement as a treatment for severe KLHL40-associated nemaline myopathy.

## INTRODUCTION

Nemaline myopathy (NM) is a clinically and genetically heterogeneous group of skeletal muscle disorders characterized by muscle weakness, hypotonia, and rod-like nemaline bodies within myofibers (1, 2). Disease onset and severity span a broad spectrum, ranging from mild, later-onset forms to severe congenital presentations that manifest in infancy with profound motor impairment and respiratory insufficiency; these congenital forms represent the most severe end of the disease spectrum and are associated with high perinatal mortality (3). Despite advances in defining the genetic basis of NM, current management remains largely supportive, and no disease-modifying therapies exist for patients with severe disease. Among the known genetic causes, recessive mutations in *KLHL40* lead to one of the most severe congenital NM subtypes (Nemaline myopathy 8, NEM8), characterized by profound muscle weakness and frequent early lethality, underscoring the urgent need for targeted therapeutic strategies (4–6).

*KLHL40* encodes a sarcomere-specific member of the BTB-Kelch family that functions as a substrate adaptor for Cullin3-RING E3 ubiquitin ligase and regulates sarcomere assembly through distinct substrate-specific mechanisms (7–9). As a canonical Cullin3 adaptor, KLHL40 promotes ubiquitin-dependent degradation of selected substrates in skeletal muscle, including SAR1A, to regulate skeletal muscle function, as we previously demonstrated (8). KLHL40 also stabilizes key thin filament proteins, including leiomodin 3 (LMOD3) and nebulin (NEB), thus protecting them from premature degradation and thereby supporting thin filament assembly, elongation, and maintenance (9). These complementary activities place KLHL40 at a central point in skeletal muscle proteostasis, balancing selective protein turnover with sarcomere assembly and function. Consequently, loss of KLHL40 destabilizes essential sarcomeric proteins, impairs myofibril assembly, and disrupts sarcomere organization, resulting in nemaline body formation, severe muscle weakness, and early lethality (4, 9). Although previous studies have established an essential role for KLHL40 in skeletal muscle development and sarcomere formation, it remains unknown whether restoring KLHL40 expression can prevent or reverse pathological defects *in vivo*.

Gene replacement has emerged as a promising therapeutic strategy for monogenic neuromuscular disorders caused by loss-of-function mutations (10–16). Systemic adeno-associated virus serotype 9 (AAV9)-mediated gene delivery has demonstrated robust skeletal muscle transduction and therapeutic efficacy across multiple preclinical models, including Duchenne muscular dystrophy, X-linked myotubular myopathy, and spinal muscular atrophy, and has translated into clinical benefit in several neuromuscular diseases (13, 17–22). These advances provide strong proof of concept that restoring the missing gene can improve muscle function and alter disease progression. The genetic and biological characteristics of KLHL40-associated nemaline myopathy, including its monogenic etiology, primarily muscle-restricted expression, and loss-of-function mechanism, make it an attractive candidate for gene replacement therapy. However, whether *in vivo* delivery can achieve therapeutically relevant expression throughout skeletal muscle, restore sarcomere integrity and contractile function, and improve lifespan without toxicity remains unknown.

Here, we evaluated the therapeutic potential of systemic AAV9-mediated *KLHL40* gene replacement in a mouse model of severe nemaline myopathy. Using KLHL40-deficient mice, we tested whether a single intravenous administration of an AAV9 vector expressing human *KLHL40* during the early disease state could rescue survival, improve growth and motor performance, restore skeletal muscle structure and function, and correct the underlying pathological abnormalities. AAV9-mediated *KLHL40* delivery resulted in robust, sustained therapeutic benefit, including near-complete rescue of survival, restoration of muscle morphology and sarcomere organization, improved muscle function, and no detectable treatment-related histopathological abnormalities in major organs. Together, these findings provide compelling preclinical evidence supporting AAV9-mediated KLHL40 gene replacement as a therapeutic strategy for patients with severe *KLHL40*-associated nemaline myopathy.

## RESULTS

### *Klhl40* knockout mice provide a robust preclinical model of *KLHL40* nemaline myopathy

*KLHL40* nemaline myopathy is a severe congenital muscle disease with no effective disease-modifying therapy. To establish a preclinical platform for evaluating AAV-based gene replacement, we obtained a constitutive *Klhl40* knockout (KO) line (C57BL/6NJ-*Klhl40*^em1(IMPC)J^/Mmjax, MMRRC, The Jackson Laboratory) generated by CRISPR/Cas9 editing. Two guide RNAs (5′-GGTTAGGCAGATAAAAAGGG-3′ and 5′-GGACCACAGCAGCAACCAGG-3′) introduced a 471-bp deletion spanning exon 2 (ENSMUSE00000633486) and 310 bp of flanking intronic sequence (Figure 1A), disrupting the open reading frame and predicting premature translational termination. Germline transmission was confirmed, and wild-type (WT), heterozygous (HT), and homozygous mutant (MT) offspring were distinguished by PCR genotyping, which yielded amplicons of approximately 700 bp for the WT allele and 200 bp for the mutant allele (Figure 1B). *Klhl40* mRNA was reduced by more than 95% in mutants relative to controls (Figure S1A), and western blotting confirmed complete absence of KLHL40 protein in homozygous mutants (hereafter referred to as knockout, KO), establishing the mutant allele as a functional null (Figures 1C-D, S2).

**Figure 1.**
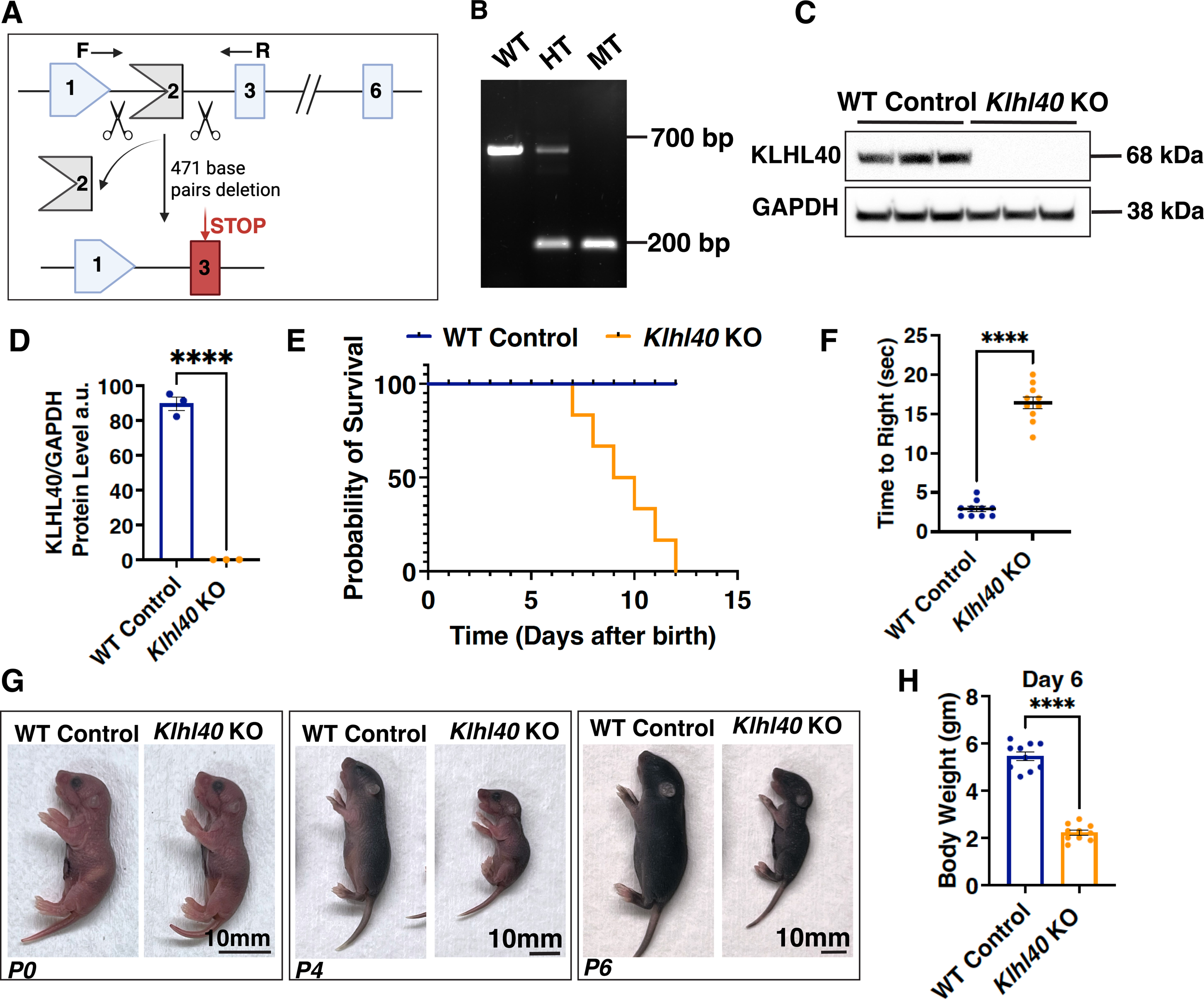
Generation and characterization of a *Klhl40* KO mouse model. (A) Schematic of the CRISPR/Cas9 targeting strategy used to generate the *Klhl40* KO mouse. Guide RNAs (scissors) flanking exon 2 direct a 471-base-pair deletion, disrupting the open reading frame. F and R indicate the forward and reverse genotyping primers. (B) Representative PCR genotyping gel of the genomic DNA distinguishing wild-type (WT), heterozygous (HT), and mutant (MT) alleles (MT allele is hereafter referred to as knockout; KO). The WT allele produces a ∼700-bp band, while the deleted KO allele produces a ∼200-bp band; heterozygotes show both bands. (C) Representative Western blot of KLHL40 protein in skeletal muscle lysates from control and *Klhl40* KO mice, confirming loss of KLHL40 protein in KO animals. GAPDH was used as a loading control. (D) Quantification of KLHL40 protein levels normalized to GAPDH in control and *Klhl40* KO muscle (n = 3 per group). Data are presented as mean ± SEM; P < 0.0001, two-tailed unpaired t-test. (E) Kaplan-Meier survival curve showing probability of survival over the first month of life in control (blue) versus *Klhl40* KO (orange) mice. *Klhl40* KO mice exhibit markedly reduced postnatal survival, with complete lethality by approximately postnatal day 14 (∼0.5 months). (F) Righting reflex assay showing time to right (seconds) in control versus *Klhl40* KO neonatal mice (n = 9-10 pups per group from three different litters). *Klhl40* KO mice show significantly prolonged time to right, indicating impaired neuromuscular function. Data are mean ± SEM; P < 0.0001, two-tailed unpaired t-test. (G) Representative whole-body images of control and *Klhl40* KO mice at postnatal day 0 (P0), day 4 (P4), and day 6 (P6), illustrating progressive growth impairment and reduced body size in KO animals. Scale bars, 10 mm. (H) Quantification of body weight (grams) in control versus *Klhl40* KO mice at postnatal day 6 (n = 8-9 pups from three different litters). *Klhl40* KO mice show significantly reduced body weight compared to controls. Data are mean ± SEM; P < 0.0001, two-tailed unpaired t-test.

*Klhl40* KO mice were born at the expected Mendelian ratio (Figure S1B), indicating that KLHL40 is dispensable for embryonic development and prenatal survival. Mutant pups appeared grossly normal at birth, with body weights indistinguishable from littermates at postnatal day 0 (P0). Between P6 and P14, however, all homozygous mutants died (Figure 1E), defining a narrow but consistent postnatal window in which KLHL40 becomes essential. By P6, KO pups showed a marked deficit in self-righting, a standard measure of axial and proximal muscle strength in neonatal rodents (23), mirroring the profound congenital hypotonia characteristic of severe KLHL40-associated nemaline myopathy in patients (Figure 1F) (4). A progressive growth deficit emerged in parallel, becoming apparent by P4 and highly significant by P6 (Figures 1G-H; \*\*\**p* < 0.001), consistent with failure to thrive.

Together, the congenital onset, progressive weakness, failure to thrive, and neonatal lethality of *Klhl40* KO mice closely recapitulate the human disease. The severity and early onset of the phenotype provide a short but well-defined therapeutic window, with survival, body-weight trajectory, and motor function serving as quantifiable endpoints for evaluating AAV-mediated *KLHL40* gene replacement.

### KLHL40 deficiency impairs postnatal myofiber growth

To determine the effect of KLHL40 loss on postnatal muscle development, we compared hindlimb morphology and myofiber composition between WT and *Klhl40* KO mice at P0 and P6. Gross muscle morphology was comparable between genotypes at birth, but by P6 the hindlimb muscles of KO mice were visibly smaller than those of WT littermates (Figure 2A-B), indicating a defect that emerges after birth rather than a failure of prenatal muscle development.

**Figure 2.**
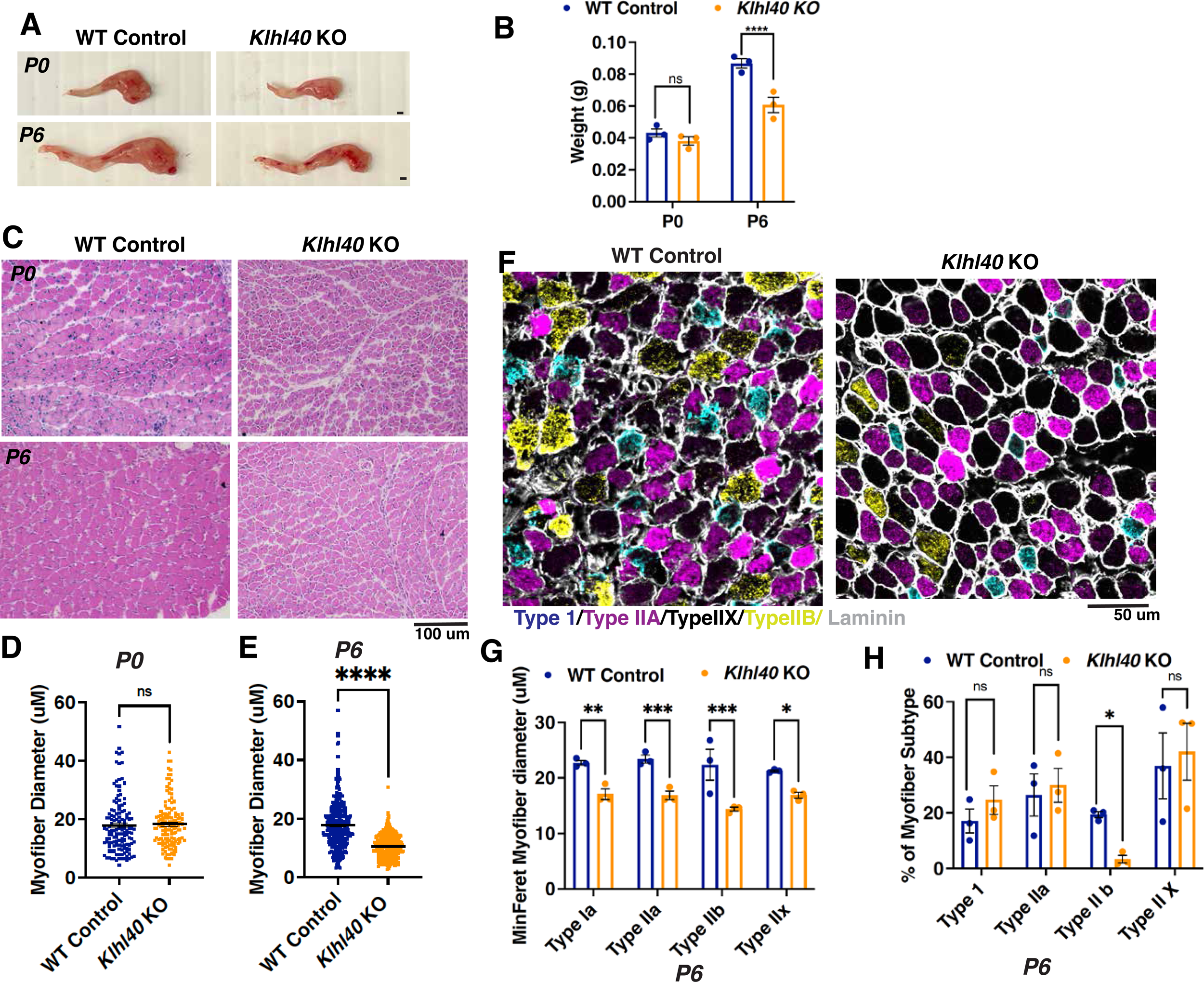
KLHL40 deficiency impairs postnatal myofiber growth. (A) Representative images of hindlimb muscle from WT control and *Klhl40* KO mice at P0 and P6, showing comparable gross morphology at birth but visibly reduced muscle size in KO mice by P6. Scale bars, 1 mm. (B) Quantification of hindlimb muscle weight (g) in WT control and *Klhl40* KO mice at P0 and P6 (n = 6-8 per group from three different litters). Muscle weight was comparable between genotypes at P0 (ns) but significantly reduced in KO mice by P6 (P < 0.0001, two-way ANOVA followed by uncorrected Fisher’s LSD test, with significance set at α = 0.05). (C) Representative hematoxylin and eosin (H&E)-stained cross-sections of hindlimb muscle from WT control and *Klhl40* KO mice at P0 and P6. Overall myofiber architecture is preserved in KO mice at both ages, with no evidence of degeneration, necrosis, or inflammatory infiltrate. Scale bar, 100 μm. (D, E) Quantification of myofiber diameter (μm) at (D) P0 and (E) P6 in WT control and *Klhl40* KO mice. Myofiber diameter is equivalent between genotypes at P0 (ns) but significantly reduced in KO mice by P6 (P < 0.0001, two-tailed, unpaired t-test) (n = 3 animals per group; each replicate is the mean of 380-546 myofibers from technical replicates). (F) Representative multiplex immunofluorescence images of hindlimb muscle cross-sections from WT control and *Klhl40* KO mice at P6, stained for myosin heavy chain isoforms to distinguish type I (blue), type IIa (magenta), type IIx (black, MyHC-I/IIa/IIb-negative), and type IIb (yellow) fibers, with laminin (white) marking fiber boundaries. Scale bar, 50 μm. (G) Quantification of minimum Feret diameter (μm) by fiber type in WT control and *Klhl40* KO muscle at P6 and (H) Quantification of fiber-type distribution (% of total myofibers) in WT control and *Klhl40* KO muscle at P6. (n = 3 animals per group; each replicate is a mean of 10-80 myofibers from technical replicates) (*P < 0.0332; **P < 0.021; ***P < 0.0002, two-way ANOVA followed by Tukey test, with significance set at α = 0.05). Data are presented as mean ± SEM.

Hematoxylin and eosin (H&E) staining showed preserved overall muscle architecture in *Klhl40* KO mice at both ages, with no evidence of degeneration, necrosis, or inflammatory infiltrate (Figure 2C). Despite the absence of overt degenerative pathology, ultrastructural analysis at P3 revealed early disruption of myofibrillar organization in *Klhl40* KO muscle compared with the well-aligned sarcomeres of WT controls (Figure S3A-B). By P6, modified Gomori trichrome staining demonstrated abnormal subsarcolemmal and intrafiber accumulations in KO myofibers consistent with nemaline pathology, which were absent from WT muscle (Figure S3C-D). Myofiber diameter was equivalent between genotypes at P0 but significantly reduced in KO mice by P6 (Figures 2D-E; \*\*\*\**p* < 0.0001), indicating that myofiber hypotrophy develops during the early postnatal period. Multiplex immunofluorescence for myosin heavy chain isoforms at P6 classified fibers as type I, IIa, IIb, or IIx (MyHC-I/IIa/IIb-negative) (Figure 2F). Minimum Feret diameter was reduced across every fiber type in KO muscle (type I, *p* < 0.021; type IIa and IIb, *p* < 0.0002; type IIx, *p* < 0.0332; Figure 2G), indicating a global rather than fiber-type-restricted growth defect. Fiber-type distribution was largely preserved, with the exception of a selective reduction in the proportion of type IIb fibers (*p* < 0.0332; Figure 2H). Loss of KLHL40 therefore produces a broad postnatal hypotrophy affecting both slow- and fast-twitch myofiber populations, with an additional more selective effect on type IIb fiber abundance.

### Systemic KLHL40 delivery prolongs survival of KLHL40-deficient mice

To test whether KLHL40 gene replacement could rescue the severe neonatal muscle weakness and lethality, we engineered a muscle-tropic AAV9 vector expressing human KLHL40 cDNA under a muscle creatine kinase promoter (AAV9-tMCK-KLHL40; hereafter AAV9-KLHL40) and administered it by retro-orbital injection at P0 (Figure 3A) (24). Three doses were tested (5 X 10¹², 1 X 10¹³, and 5 X 10¹³ vg/kg). The two lower doses extended survival only transiently, with treated KOs succumbing within the first month of life with only modestly improved survival relative to untreated KOs, indicating that sub-threshold vector exposure is insufficient to prevent lethality. In contrast, the highest dose (5 X 10¹³ vg/kg) markedly rescued lethality, with more than 80% of treated KOs surviving throughout an 8-month observation period with survival not significantly different from WT (Figure 3B), demonstrating a clear dose-dependent threshold for therapeutic efficacy. All subsequent experiments were performed with 5 X 10¹³ vg/kg.

**Figure 3.**
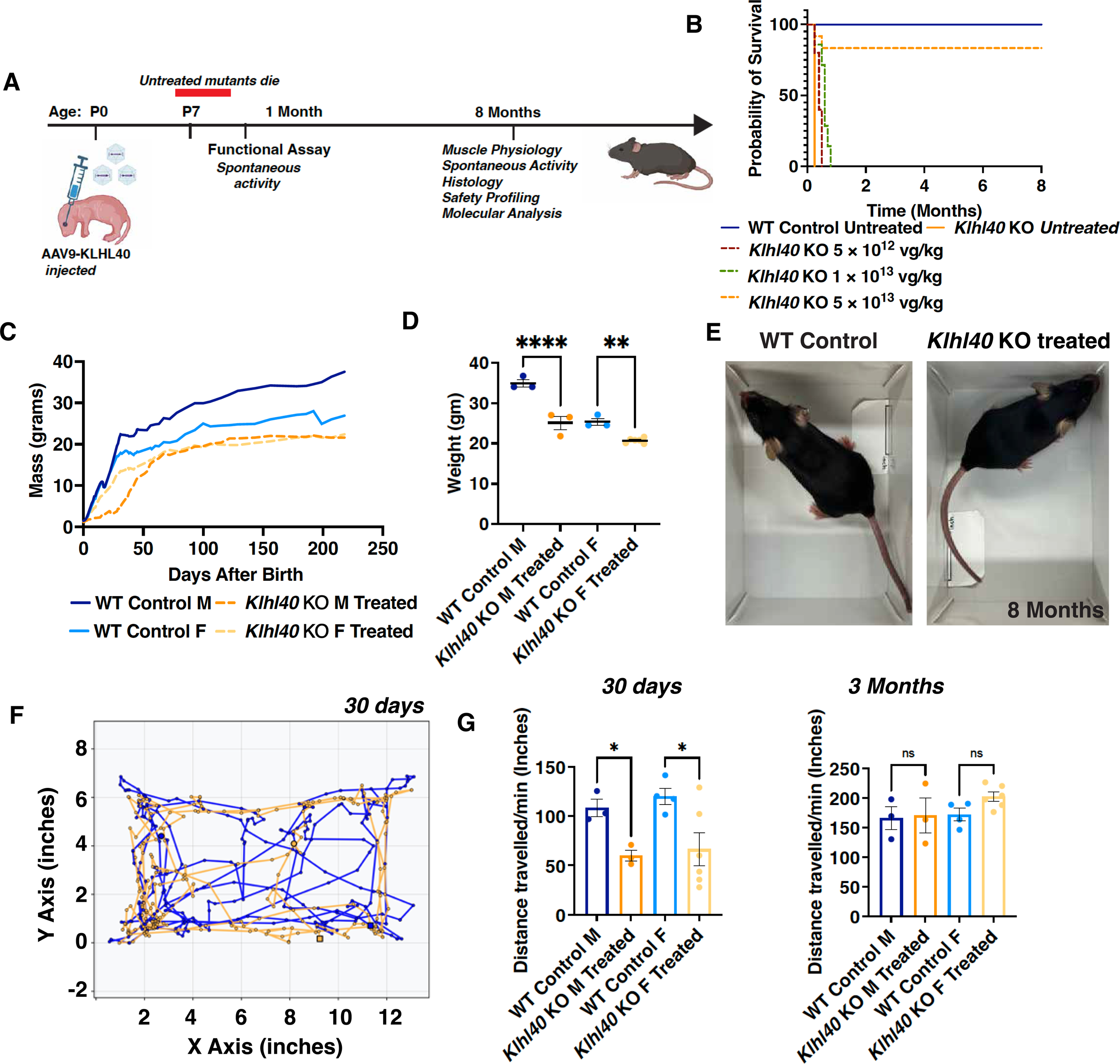
Neonatal AAV9-KLHL40 gene therapy rescues survival and confers sustained, dose-dependent functional benefit in *Klhl40* KO mice. (A) Experimental timeline. Neonatal (P0) mice received a single systemic injection of AAV9-KLHL40. Untreated *Klhl40* KO mice die between P6 and P14. Spontaneous activity was analyzed from 1 month of age, and muscle physiology, spontaneous activity, histology, safety profiling, and molecular analyses were performed at 8 months. (B) Kaplan-Meier survival curve showing probability of survival over 8 months for control untreated mice, untreated *Klhl40* KO mice, and *Klhl40* KO mice treated with AAV9-KLHL40 at three doses (5 X 10¹², 1 X 10¹³, and 5 X 10¹³ vg/kg). Untreated *Klhl40* KO mice die between P6 and P14, while AAV9-KLHL40 treatment rescues survival in a dose-dependent manner, with the highest dose (5 X 10¹³ vg/kg) achieving survival comparable to controls through 8 months (n=10 in each group, 3-8 different litters). All subsequent analyses were performed using AAV-KLHL40-treated KO mice at 5 X 10¹³ vg/kg. (C) Body mass gain trajectory over time (days after birth) for control male, control female, AAV-KLHL40-treated *Klhl40* KO male, and AAV-KLHL40-treated *Klhl40* KO female mice. (D) Body weight in control versus AAV-KLHL40-treated *Klhl40* KO mice, separated by sex at the 8-month endpoint. AAV-KLHL40-treated *Klhl40* KO mice showed steady, consistent weight gain throughout the study, though the final body weight remained significantly below sex-matched controls (Data are shown as mean ± SEM, male: P < 0.0001; female: P < 0.01, two-tailed unpaired t-test) (n = 3-6 animals in each group from three biological replicates). (E) Representative images of WT control and AAV-KLHL40-treated *Klhl40* KO mice at 8 months of age (males), illustrating overall body size and posture. (F) Representative open-field activity traces at 30 days post-treatment for control (blue) and AAV-KLHL40-treated *Klhl40* KO (orange) mice, plotted as X-Y position over time. (G) Quantification of distance traveled per minute during open-field testing at 30 days of age. AAV-KLHL40-treated *Klhl40* KO mice demonstrated measurable locomotor activity by 30 days of age, an early sign of functional rescue, though activity levels remained significantly reduced relative to sex-matched controls at this time point (Data are shown as mean ± SEM; P < 0.05, two-tailed unpaired t-test) (n=3-6 animals in each group from three biological replicates). (H) Quantification of distance traveled per minute during open-field testing at 3 months of age. No significant differences in activity were observed between control and AAV-KLHL40-treated *Klhl40* KO mice of either sex (Data are shown as mean ± SEM, ns, not significant, two-tailed unpaired t-test), indicating recovery of locomotor activity over time (n = 3-6 animals in each group from three biological replicates).

Treated *Klhl40* KO mice demonstrated steady, consistent weight gain throughout the study, reaching approximately 74% of WT body weight in males and 80% in females by the 8-month endpoint (Figure 3C-D), a substantial improvement over the untreated phenotype, in which KO mice do not survive past 1-2 weeks of life (Figure 1). Consistent with this substantial recovery of body weight, treated mice displayed grossly normal posture, coat condition, and overall body habitus at 8 months, closely resembling WT controls in general appearance (Figure 3E). Motor recovery followed a similar trajectory of progressive improvement: by 30 days, treated mice had already achieved roughly half of WT spontaneous activity levels, representing an early partial functional rescue (Figure 3F-G), and by 3 months, locomotor activity was statistically indistinguishable from WT, indicating full recovery of motor function (Figure 3H). Therapeutic benefit of AAV9-KLHL40 was observed in both male and female mice, indicating that treatment efficacy was not restricted to either sex. Taken together, these findings indicate that a single systemic neonatal dose of AAV9-KLHL40 is sufficient to prevent lethality and drive a progressive, substantial recovery of growth and motor function.

### AAV9-KLHL40 treatment restores skeletal muscle mass and contractile function

Having established that AAV9-KLHL40 prevents lethality and drives recovery of growth and locomotor activity, we next asked whether treatment also restored muscle mass and contractile function at the level of individual muscles. Gross morphological examination of the gastrocnemius, quadriceps, extensor digitorum longus (EDL), and soleus revealed no overt differences in size, shape, or color between treated KO and WT muscles (Figure 4A), suggesting that treatment prevents the gross hypotrophy characteristic of untreated *Klhl40* KO disease. Consistent with this, muscle weights across all four muscles were indistinguishable between treated KOs and sex-matched WT controls (Figure 4B), with this recovery evident across muscles spanning a range of fiber-type composition and function, from the predominantly fast-twitch EDL to the slower, more oxidative soleus, indicating that the therapeutic benefit is not restricted to a specific muscle group or fiber-type profile. This normalization of muscle mass was observed for both sexes, consistent with the therapeutic benefit observed in male and female mice (Figure 3).

**Figure 4.**
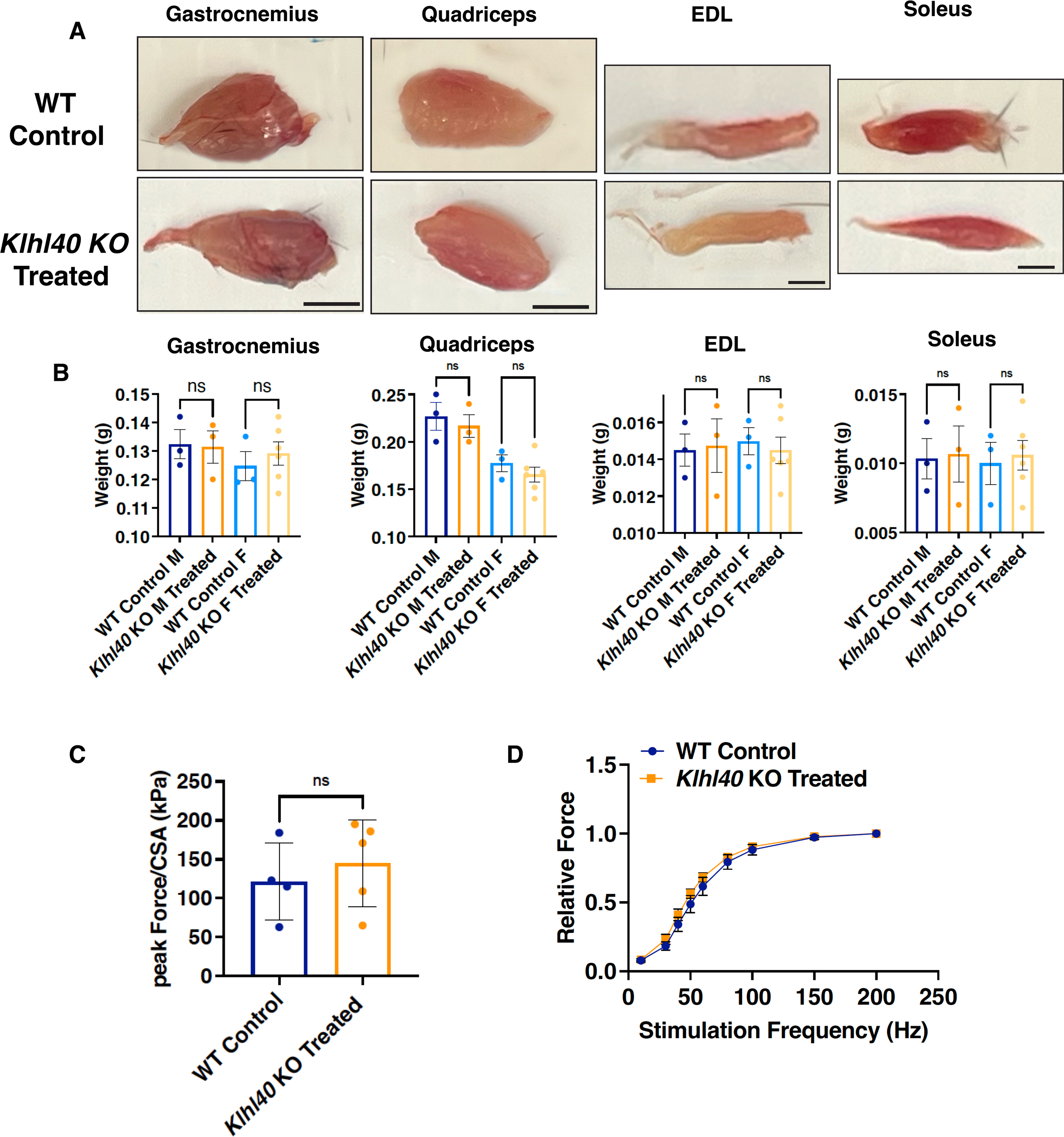
AAV-KLHL40 treatment restores skeletal muscle mass and contractile function in *Klhl40* KO mice. (A) Representative gross morphology of gastrocnemius, quadriceps, extensor digitorum longus (EDL), and soleus muscles dissected from wild-type control and AAV-KLHL40-treated *Klhl40* KO mice at 8 months after treatment, showing comparable size and shape between genotypes. Scale bars: 5 mm (gastrocnemius, quadriceps) and 2.5 mm (EDL, soleus). (B) Quantification of muscle wet weight for gastrocnemius, quadriceps, EDL, and soleus muscles, separated by sex (M, male; F, female). No significant differences in muscle mass were observed between control and AAV-KLHL40-treated *Klhl40* KO animals for either sex in any muscle group (ns, not significant). Data are shown as mean ± SEM; ns, not significant; two-tailed unpaired t-test (n = 3-6 animals in each group). (C) Peak isometric force normalized to physiological cross-sectional area (peak Force/pCSA) in control versus AAV-KLHL40-treated *Klhl40* KO muscle, showing no significant difference between groups (ns). Data are shown as mean ± SEM; two-tailed unpaired t-test (n = 3-6 animals in each group from three biological replicates). (D) Force-frequency relationship showing relative force generated across a range of stimulation frequencies (10-200 Hz) in control (blue) versus AAV-KLHL40-treated *Klhl40* KO (orange) soleus muscle, demonstrating comparable frequency-dependent force generation between groups. Force-frequency curves were compared using nonlinear regression (sigmoidal 4-parameter fit) followed by an extra sum-of-squares F test (data are plotted as mean ± SD). No significant difference was observed in the force-frequency relationship between control and AAV-KLHL40-treated *Klhl40* KO muscle (n = 3-6 animals in each group from three biological replicates).

Functional recovery in treated mice also closely paralleled the structural rescue. Peak force normalized to physiological cross-sectional area (peak force/pCSA), which reflects force-generating capacity relative to muscle size, was comparable between treated KO and WT muscles (Figure 4C), indicating restoration of specific force following AAV9-KLHL40 treatment. Force-frequency curves further supported this conclusion with treated KO and WT muscles exhibiting similar frequency-dependent increases in force across the stimulation range tested and reaching comparable maximal force (Figure 4D). These findings demonstrate that AAV9-KLHL40 treatment restores skeletal muscle’s capacity to generate force across increasing stimulation frequencies. Together, these physiological data demonstrate that normalization of muscle mass is accompanied by restoration of force-generating capacity, indicating that AAV9-KLHL40 treatment rescues both structural and functional deficits. Taken together with the growth and locomotor findings (Figure 3), these results demonstrate that a single neonatal dose of AAV9-KLHL40 drives durable, functionally meaningful rescue of skeletal muscle across multiple structural and physiological domains.

### AAV9-KLHL40 gene replacement restores sarcomere architecture and corrects the histopathological features of KLHL40 deficiency

To determine whether the functional improvement following AAV9-KLHL40 treatment was accompanied by correction of the underlying muscle pathology, we examined skeletal muscle by histological, ultrastructural, and immunofluorescence analyses. H&E staining showed that overall muscle architecture in AAV9-KLHL40-treated KOs was comparable to WT muscle (Figure 5A). At the ultrastructural level, untreated KO muscle showed severe myofibrillar disorganization as early as P3 (Figure S3), with disrupted sarcomere alignment, misaligned Z-discs, and abundant electron-dense nemaline bodies, a defining feature of the human disease. In contrast, muscle from treated KOs examined at 8 months showed well-organized myofibrils with regularly spaced sarcomeres, aligned Z-discs, and normal thick- and thin-filament organization. Nemaline bodies were not detected by electron microscopy in treated muscle (Figure 5B-C). Consistent with these findings, sarcomere height and length were restored in treated KOs and were comparable to WT (Figure 5D).

**Figure 5.**
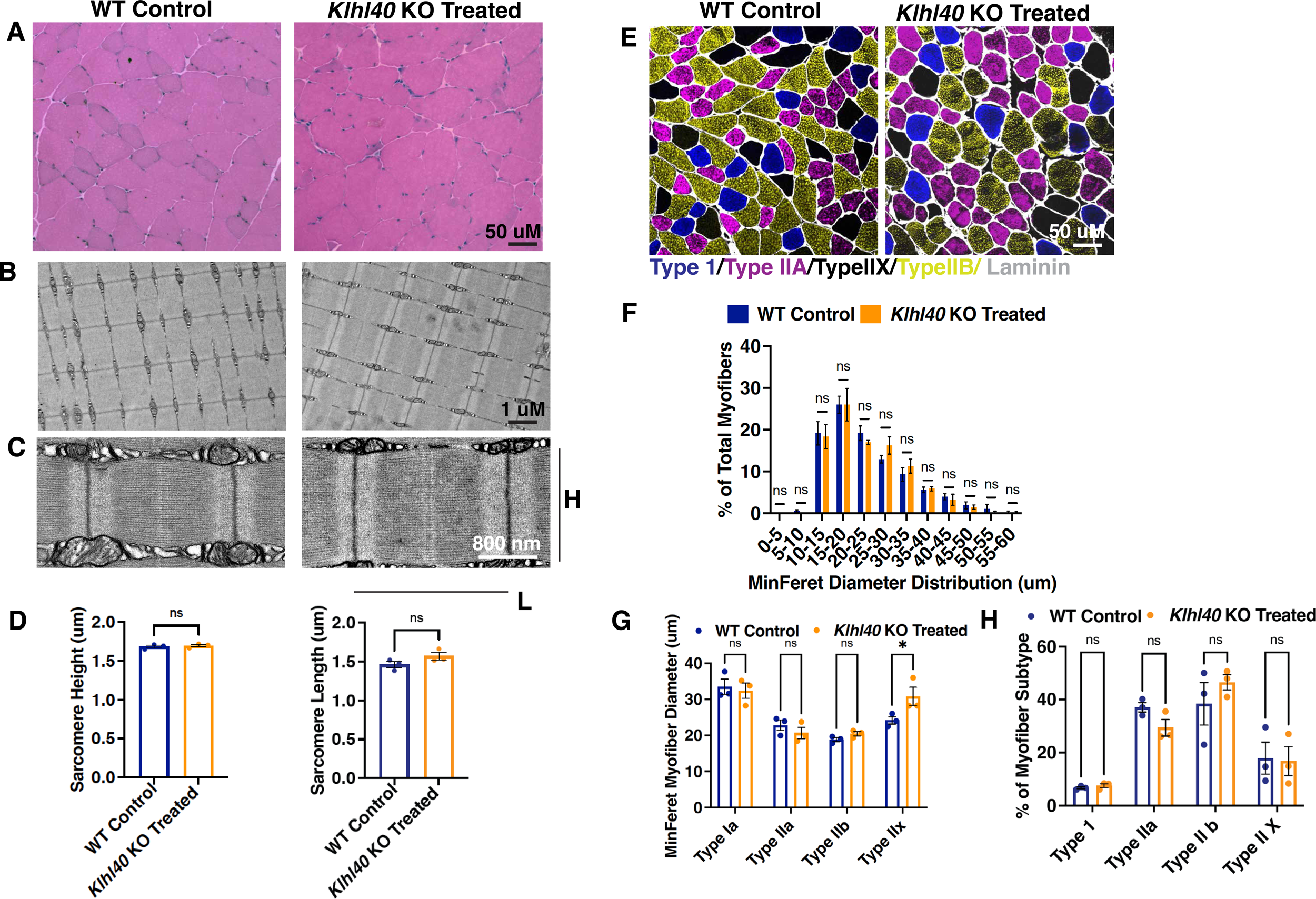
AAV9-KLHL40 treatment restores normal muscle histology, sarcomeric ultrastructure, and fiber-type composition in *Klhl40* KO mice. (A) Representative hematoxylin and eosin (H&E)-stained cross-sections of skeletal muscle from wild-type control and AAV-KLHL40-treated *Klhl40* KO mice, showing comparable fiber morphology and organization (8 months). Scale bar: 50 µm. (B) Representative transmission electron microscopy (TEM) images of longitudinal muscle sections from wild-type control and AAV-KLHL40-treated *Klhl40* KO mice, showing normal myofibrillar organization. Scale bar: 1 µm. (C) Higher-magnification TEM images of sarcomeric ultrastructure in wild-type control and AAV-KLHL40-treated *Klhl40* KO muscle. H, Sarcomere height; L, Sarcomere length. Scale bar: 800 nm. (D) Quantification of sarcomere height (µm) and sarcomere length (µm) from TEM images in control versus AAV-KLHL40-treated *Klhl40* KO muscle. No significant differences were observed (ns). Data points are from the mean of technical replicates (75-100 sarcomeres in each replicate) from three different groups of biological replicates (Data are shown as mean ± SEM; ns, not significant, two-tailed unpaired t-test). (E) Representative immunofluorescence images of muscle cross-sections from wild-type control and AAV-KLHL40-treated *Klhl40* KO mice, stained for myosin heavy chain isoforms to identify fiber type (Type I, blue; Type IIA, magenta; Type IIX, black; Type IIB, yellow) and laminin (gray) to delineate fiber boundaries. Scale bar: 50 µm. (F) Quantification of myofiber minimum Feret diameter distribution (5-µm bins) as a percentage of total myofibers in control versus AAV-KLHL40-treated *Klhl40* KO muscle. No significant differences were observed across any size bin (ns). Data are presented as mean ± SEM, with individual data points shown; two-way ANOVA followed by Tukey’s test, with significance set at α = 0.05 (n = 120-150 myofibers per replicate, 3 biological replicates per group). (G) Quantification of minimum Feret myofiber diameter (µm) by fiber subtype (Type I, IIa, IIb, IIx) in wild-type control versus AAV-KLHL40-treated *Klhl40* KO muscle. No significant differences were observed for Type I, IIa, or IIb fibers; Type IIx fiber diameter was significantly greater in AAV-KLHL40-treated *Klhl40* KO muscle (P < 0.05). Data are presented as mean ± SEM with individual data points shown, two-way ANOVA followed by Tukey test, with significance set at α = 0.05 (n = 120-150 myofibers in each replicate, 3 biological replicates in each group). (H) Quantification of the percentage of each myofiber subtype (Type 1, IIa, IIb, IIx) in wild-type control versus AAV-KLHL40-treated *Klhl40* KO muscle. No significant differences in fiber-type composition were observed (ns). Data are presented as mean ± SEM with individual data points shown, two-way ANOVA followed by Tukey test, with significance set at α = 0.05 (n = 120-150 myofibers in each replicate, 3 biological replicates in each group).

Multiplex immunofluorescence was used to assess myofiber size and fiber-type composition (Figure 5E). Minimum Feret diameter distributions in treated KOs closely overlapped those of WT across the range of fiber sizes examined (Figure 5F). Fiber-type-specific analysis showed similar fiber diameters between treated KOs and WT for Type I, IIa, and IIb fibers, whereas Type IIx fibers were modestly but significantly larger in treated KOs (Figure 5G). The relative proportions of Type I, IIa, IIb, and IIx fibers were comparable between treated KOs and WT (Figure 5H), indicating preservation of normal fiber-type composition.

Together, these findings show that a single neonatal dose of AAV9-KLHL40 provides sustained correction of the skeletal muscle pathology associated with KLHL40 deficiency. Treatment restored myofibrillar and sarcomeric organization, prevented nemaline body formation, and normalized myofiber size and fiber-type composition.

### Systemic AAV9-KLHL40 administration achieves preferential muscle transgene expression

To define the tissue distribution of the therapeutic vector and determine whether systemic AAV9 delivery produced sustained KLHL40 expression in skeletal muscle, we quantified vector genome copy number, *KLHL40* transcript levels, and protein expression across skeletal muscle and major organs. Vector genomes were present in every tissue examined, with 4-7 copies per diploid genome across skeletal muscles (EDL, soleus, tibialis anterior [TA], and quadriceps). As expected for systemic AAV9 delivery, biodistribution was highest in liver (∼19 copies/diploid genome), with lower levels in kidney, heart, brain, and lung (∼3-5 copies/diploid genome) (Figure 6A).

**Figure 6.**
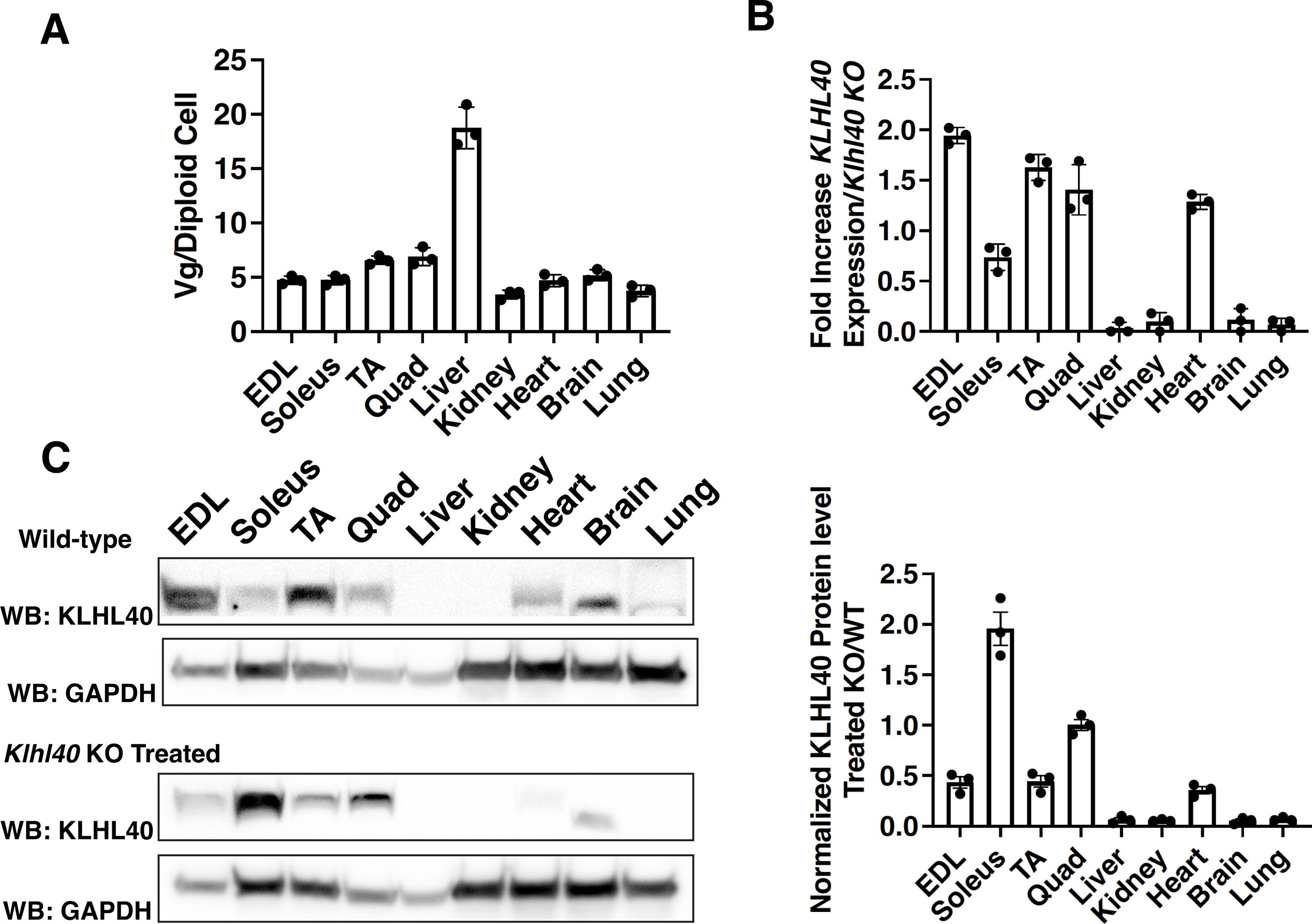
Biodistribution, transcript, and protein expression of KLHL40 following AAV-KLHL40 gene therapy. (A) Vector genome biodistribution, expressed as vector genomes per diploid genome (Vg/diploid genome), across extensor digitorum longus (EDL), soleus, tibialis anterior (TA), quadriceps (Quad), liver, kidney, heart, brain, and lung following systemic AAV-KLHL40 administration. (B) Normalized Fold increase in human *KLHL40* mRNA expression in AAV-KLHL40-treated *Klhl40* KO tissues relative to untreated *Klhl40* KO tissues, measured by qRT-PCR, across the same panel of tissues. (C) Representative western blots for KLHL40 protein in wild-type (top) and AAV-KLHL40-treated *Klhl40* KO (bottom) tissues, with GAPDH as a loading control. (D) Quantification of KLHL40 protein levels in AAV-KLHL40-treated *Klhl40* KO tissues normalized to GAPDH and expressed relative to wild-type levels (treated KO/WT). Abbreviations: EDL, extensor digitorum longus; TA, tibialis anterior; Quad, quadriceps. Data are presented as mean ± SEM; individual data points represent biological replicates (n = 3 per group).

**Figure 7.**
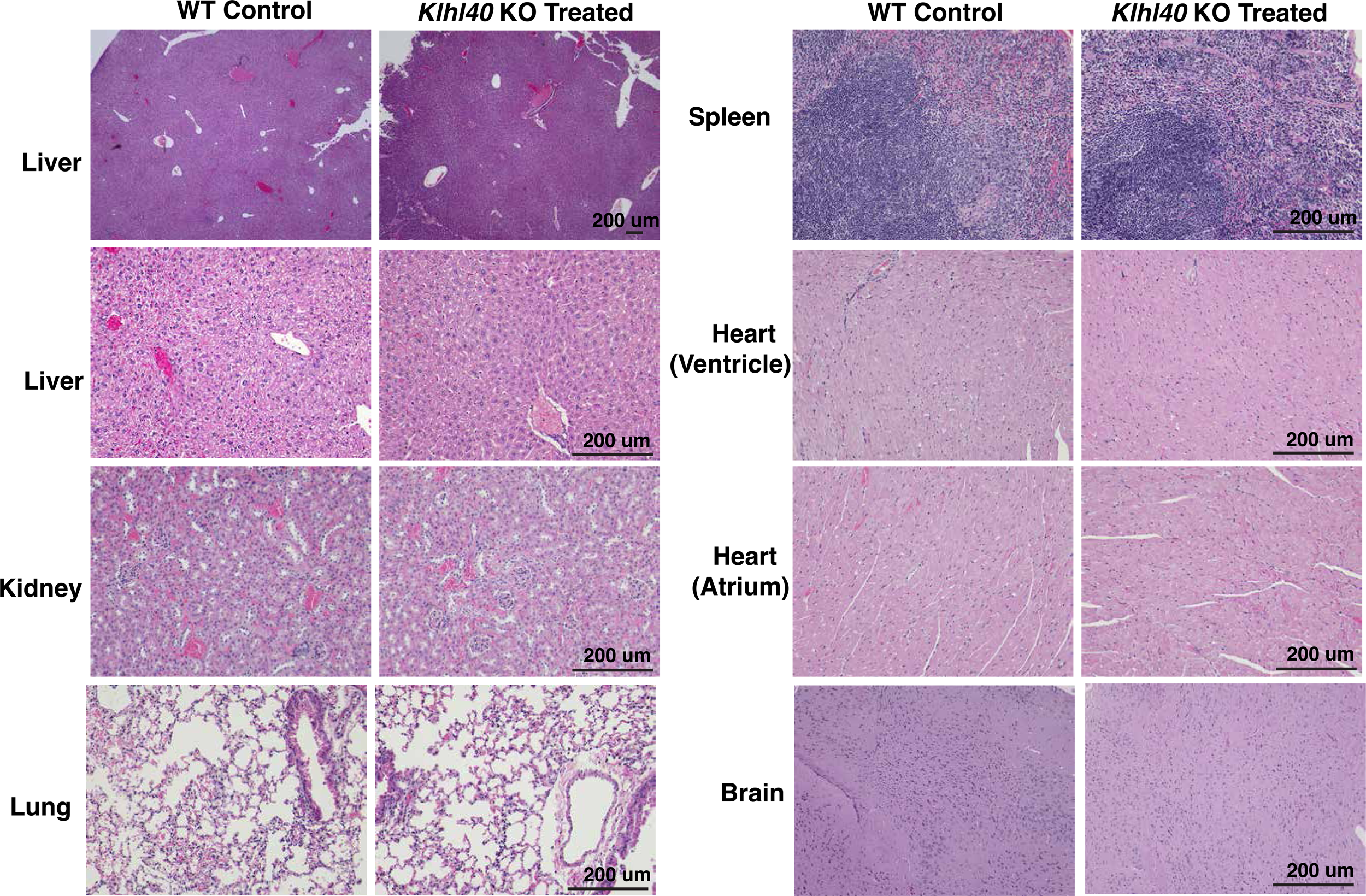
Histological analysis of major organs following AAV-KLHL40 treatment. Representative hematoxylin and eosin (H&E)-stained sections of liver (low and high magnification), kidney, lung, spleen, heart (ventricle and atrium), and brain from WT and AAV-KLHL40-treated *Klhl40* KO mice. No overt histopathological abnormalities were observed in AAV-KLHL40-treated KO tissues compared with WT controls, indicating no detectable treatment-associated histopathology in the organs examined. Scale bars, 200 µm (n = 3 animals per group).

Despite this broad distribution, transgene expression was predominantly observed in skeletal muscle, with lower expression in the heart. *KLHL40* mRNA was highest in EDL, TA, and quadriceps, lower in soleus, and detectable but comparatively modest in heart; liver, kidney, brain, and lung showed minimal transcript despite substantial vector genome presence (Figure 6B). Western blotting mirrored this pattern: KLHL40 protein was readily detected in EDL, soleus, TA, and quadriceps, present in heart, and essentially undetectable in liver, kidney, or lung (Figure 6C). This expression profile closely matched the endogenous tissue distribution of KLHL40 in WT mice, confirming that systemic AAV9-KLHL40 delivery achieves preferential transgene expression in the disease-relevant tissue while sparing non-muscle organs.

### Systemic AAV9-KLHL40 administration does not induce histopathological abnormalities in major organs

To evaluate systemic safety, we examined liver, kidney, lung, spleen, heart (atria and ventricles), and brain from long-term surviving KOs 8 months after a single neonatal dose of AAV9-KLHL40 (5 X 10¹³ vg/kg), comparing H&E-stained sections with age-matched WT controls. Tissue architecture was preserved in every organ examined: liver parenchyma, renal glomeruli and tubules, pulmonary alveolar structure, splenic white and red pulp, myocardial fibers, and brain parenchyma were all histologically unremarkable in treated animals. No inflammatory infiltrate, cellular degeneration, necrosis, fibrosis, or vascular lesion was identified in any tissue. Thus, the therapeutically effective dose of AAV9-KLHL40 that achieves complete phenotypic rescue in *Klhl40* KO mice was not associated with detectable off-target histopathological abnormalities after 8 months of follow-up.

## Discussion

Nemaline myopathy (NM) comprises a genetically heterogeneous group of congenital myopathies for which treatment remains largely supportive (2). However, substantial progress in disease modeling and preclinical therapeutic development has demonstrated that several features of NM are amenable to therapeutic intervention. Experimental approaches have included pharmacological enhancement of muscle contractility, modulation of disease-associated proteins, exon skipping, shRNA, and the development of genetically and physiologically well-characterized models for therapeutic discovery; however, clinical translation remains limited (25–30). KLHL40-NM is among the most severe forms of the disease. Most affected individuals present prenatally or at birth with fetal hypokinesia, profound muscle weakness, dysphagia, and respiratory insufficiency, with early mortality common (4). Here, we show that systemic delivery of AAV9-KLHL40 shortly after birth prevents lethality in *Klhl40* KO mice and produces sustained correction of major functional and pathological features of KLHL40 deficiency. At the effective dose, treatment restored muscle mass and contractile function, corrected sarcomeric organization, prevented nemaline body formation, and supported long-term survival. These findings provide preclinical evidence that restoration of KLHL40 expression can correct the severe muscle phenotype caused by KLHL40 deficiency.

Previous studies of KLHL40 and other BTB/Kelch proteins associated with NM have established the importance of this protein family in maintaining skeletal muscle structure and function (9, 31). The *Klhl40*^em1(IMPC)J/^Mmjax used in this study contains a defined 471-bp deletion encompassing exon 2 and adjacent intronic sequence and results in complete loss of detectable KLHL40 protein. The early lethality, impaired postnatal growth, muscle weakness, and severe sarcomeric abnormalities observed in this model recapitulate key features expected from loss of KLHL40 function in patients and provide a stringent setting in which to evaluate therapeutic rescue.

The extent of structural and functional recovery following AAV9-KLHL40 treatment is particularly notable given the severity and early onset of disease in untreated mice. Untreated *Klhl40* KO muscle developed hypotrophy, marked myofibrillar disruption, and nemaline bodies within the first postnatal days, whereas treated animals examined at 8 months showed normal myofiber size, organized myofibrils, normal sarcomere dimensions, and no detectable nemaline bodies. These structural improvements were accompanied by normalization of muscle-specific force and force-frequency relationships. Thus, restoring KLHL40 expression does more than prolong survival; it enables skeletal muscle to establish and maintain functional sarcomeric architecture. This finding is consistent with the established role of KLHL40 in sarcomere integrity and in regulating the stability of proteins required for thin-filament structure and function (8, 9).

The response across the three vector doses also identifies a clear efficacy threshold in this rapidly progressive model. Doses of 5 X 10^12^ and 1 X 10^13^ vg/kg provided only transient benefit, whereas 5 X 10^13^ vg/kg prevented early lethality and resulted in durable phenotypic rescue. Therapeutic benefit was consistent across male and female mice. The requirement for early systemic treatment is relevant to the natural history of NEM8, in which disease manifestations frequently begin prenatally, and severe disease is often evident at birth (6). The effective dose is also within the range previously evaluated for systemic AAV9 delivery in large-animal studies. Intravenous administration of an AAV9 vector at 3 X 10^13^ and 8 X 10^13^ vg/kg was well tolerated in juvenile nonhuman primates followed for up to 6 months, without significant biochemical, clinical, or histopathological toxicity (32). Although differences in vector genome, transgene, promoter, manufacturing, and underlying disease preclude direct comparison between programs, these studies provide useful safety context for the dose evaluated here. Engineered muscle-tropic capsids may provide an additional opportunity to increase skeletal muscle transduction while reducing the vector dose and off-target liver exposure required for therapeutic efficacy (33–35). MyoAAV or AAVmyo variants have shown substantially greater muscle transduction than AAV9 in mice and efficient muscle targeting in nonhuman primates; however, whether these preclinical advantages translate into improved efficacy and safety in patients remains to be established.

Safety remains an important consideration for systemic muscle-directed AAV therapy. High systemic vector exposures have been associated with hepatotoxicity, complement activation, and thrombotic microangiopathy (36, 37). Experience with gene replacement for X-linked myotubular myopathy (XLMTM) illustrates the complexity of interpreting safety findings across systemic AAV programs. In the ASPIRO study of AAV8-MTM1, four treated participants died with cholestatic liver failure, including three of 17 participants receiving 3.5 X 10^14^ vg/kg and one of seven receiving 1.3 X 10^14^ vg/kg (38). Importantly, MTM1 deficiency itself is associated with hepatobiliary abnormalities, suggesting that the severe hepatic events observed in ASPIRO may reflect an interaction among underlying disease biology, vector exposure, and other patient-specific factors rather than a toxicity that can be generalized across AAV products (39, 40). The therapeutic dose used in the present study, 5 X 10^13^ vg/kg, is substantially lower than those used in ASPIRO; nevertheless, the safety profile of each systemic AAV product must be established in the context of its specific vector, transgene, dose, and underlying disease.

In the present study, we identified no overt histopathological abnormalities in the organs examined at 8 months. Despite broad vector biodistribution and high vector genome levels in the liver, KLHL40 protein expression was predominantly restricted to skeletal muscle, consistent with transcriptional targeting by the muscle-specific creatine kinase promoter (tMCK). This separation between vector biodistribution and transgene expression may reduce ectopic KLHL40 expression in non-muscle tissues, although it does not eliminate potential capsid- or vector-related toxicity associated with uptake by these organs. The well-established use of AAV9 and the broad preclinical use of creatine kinase-based muscle-specific expression provide an important foundation for further development, while the safety of the complete AAV9-KLHL40 vector will require direct evaluation. Pre-existing anti-AAV9 immunity represents an additional consideration for systemic delivery because neutralizing antibodies can substantially reduce vector transduction and may affect patient eligibility (41).

Several additional limitations should be considered. First, treatment was administered at P0-P1, reflecting the rapidly progressive phenotype and early lethality of the *Klhl40* KO model. Nevertheless, the robust rescue achieved with postnatal administration demonstrates that the severe phenotype remains amenable to correction after birth. Determining how progressively delayed treatment affects outcome will be important for defining the therapeutic window and for approximating the timing realistically expected following postnatal genetic diagnosis. Second, although rescue was maintained through 8 months, longer-term studies will be needed to establish the durability of transgene expression and phenotypic correction. Despite these limitations, the magnitude and durability of rescue demonstrate that KLHL40 deficiency is highly amenable to gene replacement. A single systemic neonatal administration of AAV9-KLHL40 prevented early lethality and produced sustained improvements in growth, motor function, skeletal muscle mass, intrinsic muscle force, and sarcomeric organization in a severe *Klhl40* knockout model. The prevention of nemaline bodies and restoration of normal myofibrillar architecture further indicate that replacement of the primary disease gene addresses the underlying structural pathology in addition to improving downstream muscle function. Together, these findings establish a strong preclinical rationale for AAV9-mediated KLHL40 replacement and support further development of this approach toward clinical translation. Defining the therapeutic window and minimum effective dose, establishing long-term safety and durability, and developing clinically applicable measures of treatment response will be important next steps.

## MATERIALS AND METHODS

### *Klhl40* mouse line and husbandry

All animal procedures were performed in accordance with protocols approved by the Brigham and Women’s Hospital Institutional Animal Care and Use Committee (IACUC protocol 2021N000233) or the Boston Children’s Hospital Institutional Animal Care and Use Committee. Mice were maintained under specific pathogen-free conditions on a 12-h light/dark cycle at controlled temperature and humidity with ad libitum access to standard chow and water. C57BL/6NJ-*Klhl40*^em1(IMPC)J^/Mmjax mice (MMRRC Stock No. 046178-JAX; RRID: MMRRC_046178-JAX) were obtained from the Mutant Mouse Resource & Research Centers (MMRRC) at The Jackson Laboratory. The mutant allele was generated by CRISPR/Cas9-mediated genome editing using two guide RNAs (5′-GGTTAGGCAGATAAAAAGGG-3′ and 5′-GGACCACAGCAGCAACCAGG-3′) to delete a 471-bp genomic region encompassing exon 2 and adjacent intronic sequence, generating a predicted null allele. Wild-type (WT), heterozygous (Het), and homozygous knockout (KO) mice were identified by PCR genotyping of genomic DNA isolated from tail biopsies using allele-specific primers (F: 5’-CATCCACACAATGCAGCTGT-3’; R: 5’-TGAGCTCTCCTGCCTTTGAG-3’). Neonatal mice of both sexes were used for all experiments unless otherwise specified.

### AAV plasmids and virus production

KLHL40-expressing plasmid was constructed from pENN.AAV.tMCK.PI.eGFP.WPRE.bGH (Addgene plasmid No 105556) by replacing the eGFP with WT human KLHL40 cDNA (GenScript). Serotype 9 AAV vector was packaged in HEK293T cells by polyethylenimine (PEI)-mediated cotransfection of the WT KLHL40 tMCK plasmid, a Rep2Cap9 plasmid, and a pHelper plasmid. AAV9-KLHL40 was then purified using discontinuous density iodixanol (OptiPrepTM, Axis-Shield) gradient ultracentrifugation. The AAV containing iodixanol fraction was collected and diafiltered to remove iodixanol and concentrate the AAV vector. The final product was suspended in 0.001% F68 in DPBS, and viral titers were determined as described (42).

### AAV9-KLHL40 administration in neonatal mice

Neonatal mice received a single systemic injection of AAV9-KLHL40 under the control of the muscle creatine kinase promoter (tMCK). Injections were performed by the retro-orbital route at postnatal day 0 or 1 using a fine-gauge insulin syringe. The vector was diluted in sterile phosphate-buffered saline to the required dose and administered at 5 X 10¹², 1 X 10¹³, or 5 X 10¹³ vg/kg body weight in an injection volume (10-15 µL) appropriate for neonatal mice. Pups were briefly anesthetized by hypothermia before injection and were returned to their dam after recovery. Control wild-type littermates received an equivalent volume of saline. Animals were monitored regularly following injection for general health, growth, survival, and potential treatment-related adverse effects.

### Survival analysis

Mice were monitored daily during the first 4 weeks of life and weekly thereafter for survival and predefined humane endpoints over an 8-month observation period. Humane endpoints included markedly reduced activity, impaired mobility or inability to access food or water, progressive deterioration in body condition, and other signs of significant distress. Animals reaching a predefined humane endpoint were euthanized and recorded as an event for survival analysis. Kaplan-Meier survival curves were generated and compared between groups using the log-rank (Mantel-Cox) test.

### Righting reflex assay

Neonatal mice were placed in a supine position, and the time to return to a prone position (righting) was recorded, with a maximum cutoff of 45 seconds. Three trials were performed per pup at 1 week of age as described (23).

### Body weight and gross morphology

Body weight was measured using a calibrated balance daily during the first 4 weeks of life and weekly thereafter. Whole-body photographs were acquired at P0, P4, and P6, and at 8 months for long-term treated cohorts, with a scale bar included in each image for calibration.

### Open-field locomotor activity

Spontaneous locomotor activity was assessed in an open-field arena for 60 seconds at 30 days and 3 months of age. Animal position was tracked using Openshot Video Editor, and distance traveled per minute was calculated.

### Tissue collection and histology

Mice were euthanized by CO inhalation in accordance with the institutionally approved animal protocol at the indicated time points. Hindlimb muscles (gastrocnemius, quadriceps, EDL, soleus) were dissected, weighed, and either fixed in formalin for paraffin sectioning or flash-frozen in liquid nitrogen-cooled isopentane and embedded in OCT for cryosectioning. Sections (8 μm) were stained with hematoxylin and eosin (H&E) using standard protocols and imaged on a Zeiss microscope.

### RNA isolation and RT-qPCR

Total RNA was extracted from different tissues using the RNeasy kit (Qiagen, #74104) using the manufacturer’s instructions, and cDNA was synthesized using the First Strand cDNA Synthesis Kit (Thermo Fisher Scientific, #K1621). Human and mouse KLHL40 TaqMan probes were obtained from Thermo Fisher Scientific. Quantitative PCR was performed using an ABI Prism 7900 apparatus (Applied Biosystems) as described previously (43).

### Western blotting

Skeletal muscle (and, for biodistribution studies, liver, kidney, heart, brain, lung) was homogenized in T-PER tissue extraction buffer (Thermo Fisher Scientific, #78510) containing protease inhibitors (Millipore Sigma, #04693159001). Samples were separated on 4-12% Bis-Tris plus mini gels (Thermo Fisher Scientific, #NW04120BOX) and blotted using Trans-Blot Turbo Mini 0.2 µm Nitrocellulose (Bio-Rad, #1704158). The membranes were blocked using 5% non-fat milk powder in 1X Tris-buffered saline and 0.1% TWEEN 20 (TBST) for 1 hour at room temperature and incubated with primary antibodies overnight at 4°C. The membranes were subsequently washed and incubated with polyclonal anti-rabbit or anti-mouse IgG antibody conjugated to horseradish peroxidase. Primary antibodies and associated dilutions were: KLHL40, 1:500 (Millipore Sigma, #HPA024462) and GAPDH, 1:1000 (Cell Signaling Technology, #2118S). Secondary antibodies were anti-rabbit 1:1000 (Bio-Rad, #170-6515) and anti-mouse, 1:1000 (Bio-Rad, #170-6516). Protein band quantification was performed using ImageJ.

### Multiplex immunofluorescence and fiber typing

Cryosections were fixed in 4% paraformaldehyde, blocked with 5% goat serum, and incubated with primary antibodies against MyHC-I (DSHB, #BA-D5-s), MyHC-IIa (DSHB, #SC-71-s), MyHC-IIb (DSHB, #BF-F3-c), and laminin (Millipore Sigma, #L9393), followed by fluorophore-conjugated secondary antibodies. Slides were imaged using spinning disc confocal microscopy. Fibers were classified as type I, IIa, IIb, or IIx (negative for I/IIa/IIb), and minimum Feret diameter and fiber-type proportions were quantified per animal from 120-150 fibers/section across 3 animals per group (FIJI).

### Transmission electron microscopy

Three different control and mutant mice from three different mating pairs were analyzed by electron microscopy, and sarcomeres were visualized. Samples were fixed in formaldehyde-glutaraldehyde-picric acid in cacodylate buffer overnight at 4°C, followed by osmication and uranyl acetate staining. Subsequently, tissue samples were dehydrated in a series of ethanol washes and finally embedded in Taab epon (Marivac Ltd., Nova Scotia, Canada). Ninety-five-nanometer sections were cut with a Leica Ultracut microtome, picked up on 100-mesh Formvar-coated Cu grids, and stained with 0.2% lead citrate. Sections were viewed and imaged by JEOL 1200EX Transmission Electron Microscope (Electron Microscopy Core, Harvard Medical School). Sarcomere length was measured across all samples and found to be consistent within replicates in each genotype, indicating comparable contractile states at the time of fixation. The maximal width of a structure is inferred when the section passes through the central longitudinal plane rather than a tangential or oblique plane. For sarcomeres or myofibrils, this is identified by the simultaneous visualization of continuous and well-aligned structural landmarks, including distinct Z-lines, A-bands, I-bands, and consistent filament organization. Therefore, measurements were restricted to regions displaying the most complete and symmetric ultrastructural organization. Moreover, consistent measurements across multiple samples and independent sections further supported that these dimensions represent near-maximal structural width. While chemical fixation cannot guarantee identical contractile states, the uniformity of sarcomere lengths across samples provides empirical evidence that the fibers were in a comparable state.

### *Ex vivo* muscle physiology

We assessed skeletal muscle contractile function using established procedures (44, 45). For *ex vivo* measurements, we isolated soleus muscles from anesthetized mice and mounted them between a force transducer and a fixed post in a bicarbonate buffer equilibrated with 95% O_2_, 5% CO_2_ and maintained at 35°C. Muscles were adjusted to the length producing maximal force and stimulated with supramaximal 200-µs square-wave pulses. Twitch force was measured using single pulses, and tetanic contractions were elicited over a range of stimulation frequencies (10-200 Hz) using 400-ms trains for soleus. Force-frequency relationships were generated from the resulting contractions. Specific force was calculated by normalizing absolute force to physiological cross-sectional area (pCSA).

### Vector genome quantification

Genomic DNA was extracted from mouse tissues using the GenElute Mammalian Genomic DNA Miniprep kit (Millipore Sigma, #G1N70). A standard curve was generated using pcDNA-KLHL40 plasmid. qPCR was performed using TaqMan Universal Master Mix II (Thermo Fisher Scientific, #4440038) and targeting the SV40 polyadenylation sequence (forward primer 5′-TGGTGGTGCAAATCAAAGAACT-3′; reverse primer: 5′-AACACTTCCGTACAGGCCTAGAA-3′; probe: 5′-6FAM-CTCAGTGGATGTTGCCTT-MGB-3′). We recorded data as vector copies per 100 ng of DNA. We calculated the number of diploid genome equivalents in each DNA sample assuming a diploid genome size of 5.4 X 10^9^ bp and an average molecular mass of 650 Da per base pair. Vector genome copies determined from the qPCR standard curve were then divided by the corresponding number of diploid genome equivalents as described previously (46, 47).

### Histopathology

Liver, kidney, lung, spleen, heart (atria and ventricles), and brain were collected from long-term surviving treated KO mice and age-matched WT controls at 8 months, fixed in formalin, paraffin-embedded, sectioned, and stained with H&E (Rodent Histopathology Core, Harvard Medical School).

### Quantification and statistical analysis

Statistical analyses were performed using GraphPad Prism (10.3.1). Data are presented as mean ± SEM unless otherwise noted (force-frequency curves are presented as mean ± SD). Comparisons between two groups were made using two-tailed unpaired Student’s t-tests. Comparisons across multiple groups or factors were made using two-way ANOVA followed by Tukey’s or uncorrected Fisher’s LSD post hoc test, as indicated in figure legends, with significance set at α = 0.05. Force-frequency relationships were compared using nonlinear regression (sigmoidal four-parameter fit) followed by an extra sum-of-squares F test. Sample sizes (n) reflect the number of biological replicates (individual animals); where technical replicates (e.g., multiple myofibers or sarcomeres per animal) were averaged to generate a single biological data point, this is noted in the corresponding figure legend. Statistical significance is denoted in figure legends.

## RESOURCE AVAILABILITY

### Lead contact

Requests for resources and reagents to the corresponding author (Vandana Gupta).

### Materials availability

The AAV9-tMCK-KLHL40 vector plasmid and construct generated in this study will be made available upon request, subject to execution of a Material Transfer Agreement (MTA) with Brigham and Women’s Hospital.

### Data and code availability

This paper does not report original code. Any additional information related to the data reported in this paper is available from the corresponding author upon reasonable request.

## ACKNOWLEDGEMENTS

This work was supported by A Foundation Building Strength (AFBS) (to V.A.G.). V.A.G. is also supported by the Muscular Dystrophy Association (MDA). We thank Jennifer Casey and Sara Coleman for assistance with mouse colony management; the Harvard Medical School MicRoN Core for microscopy assistance; and the Harvard Medical School Electron Microscopy Core for expert assistance with sample preparation. We thank the Developmental Studies Hybridoma Bank (DSHB) for providing antibodies. Finally, we are deeply grateful to all individuals and families affected by nemaline myopathy for their continued support for, and contributions to, research in this field.

## AUTHORS CONTRIBUTIONS

G.R.T.: Investigation, Formal analysis, Writing; review & editing. J.M.: Investigation. J.W.: Methodology, Investigation. V.A.G.: Methodology, Formal analysis, Funding acquisition, Writing; original draft.

## DECLARATION OF INTERESTS

The authors declare no financial conflicts of interest or other competing interests that could have influenced the conduct, analysis, or interpretation of this research.

## FIGURE LEGENDS

**Figure S1. Validation of Klhl40 mRNA depletion and Mendelian inheritance ratios in *Klhl40* KO mice.** (A) Quantitative RT-PCR analysis of *Klhl40* mRNA levels, normalized to *Gapdh*, in quadriceps skeletal muscle from control and *Klhl40* KO mice, confirming near-complete loss of *Klhl40* transcript in KO animals. Data are presented as mean ± SEM; P < 0.0001, two-tailed unpaired t-test (n = 5 biological replicates in each group). (B) Genotype distribution of offspring from *Klhl40* heterozygous incrosses, shown as the percentage of wild-type (+/+), heterozygous (+/-), and KO (-/-) pups per litter across five litters (n = 6-10 in each litter). Observed ratios are consistent with the expected Mendelian distribution, indicating that *Klhl40* deletion does not cause embryonic or perinatal lethality prior to genotyping.

**Figure S2. Uncropped Western blots corresponding to Figure 1C**. Full-length, unprocessed Western blot images showing KLHL40 (top) and GAPDH (bottom) in skeletal muscle lysates from control (n = 3 biological replicates) and *Klhl40* KO (n = 3 biological replicates) mice. Molecular weight markers (kDa) are indicated on the right of each blot.

**Figure S3. Loss of KLHL40 disrupts sarcomere organization and causes nemaline pathology in skeletal muscle.** (A-B) Representative transmission electron microscopy images of skeletal muscle from wild-type control (A) and *Klhl40* KO (B) mice at P3. WT muscle shows regularly aligned myofibrils with well-organized sarcomeres, whereas *Klhl40* KO muscle displays disrupted myofibrillar organization and sarcomere architecture. Scale bar, 500 nm. (C-D) Representative modified Gomori trichrome staining of skeletal muscle from wild-type control (C) and *Klhl40* KO (D) mice at P6. KO muscle shows abnormal trichrome-positive accumulations within myofibers consistent with nemaline pathology, which are absent from WT controls. Scale bar, 25 μm.

**Figure S4. Uncropped western blots corresponding to Figure 6**. Full-length, uncropped immunoblot images corresponding to the cropped panels shown in Figure 6. (A) WT control. (B) AAV-KLHL40-treated *Klhl40* KO. Membranes were probed for KLHL40 and GAPDH as a loading control. Molecular weight markers (kDa) are indicated.

**Supplementary Movie 1. Partial phenotypic rescue in AAV9-KLHL40-treated *Klhl40* KO mice at 1 month of age.** Representative video of AAV9-KLHL40-treated *Klhl40* KO mice compared with wild-type (WT) littermates at 1 month of age. Treated KO mice remain visibly smaller in body size than WT controls and display reduced spontaneous locomotor activity, exhibiting approximately 50% of the activity levels observed in WT mice.

**Supplementary Movie 2. Improved motor activity in AAV9-KLHL40-treated *Klhl40* KO mice at 3 months of age.** Representative video of AAV9-KLHL40-treated *Klhl40* KO mice and WT littermates at 3 months of age. Treated KO mice continue to display a smaller body size relative to WT mice; however, spontaneous locomotor activity is comparable to WT, indicating substantial functional recovery.

**Supplementary Movie 3. Sustained phenotypic rescue in AAV9-KLHL40-treated *Klhl40* KO mice at 8 months of age.** Representative video of AAV9-KLHL40-treated *Klhl40* KO mice and WT littermates at 8 months of age. Treated KO mice show substantial phenotypic recovery, with spontaneous locomotor activity comparable to WT controls.

## Notes

### Competing Interest Statement

The authors have declared no competing interest.

