## Supplementary figures and images for "KLHL40 gene replacement therapy in severe nemaline myopathy improves survival and skeletal muscle function in a preclinical mouse model"

### Figure S1

Figure S1

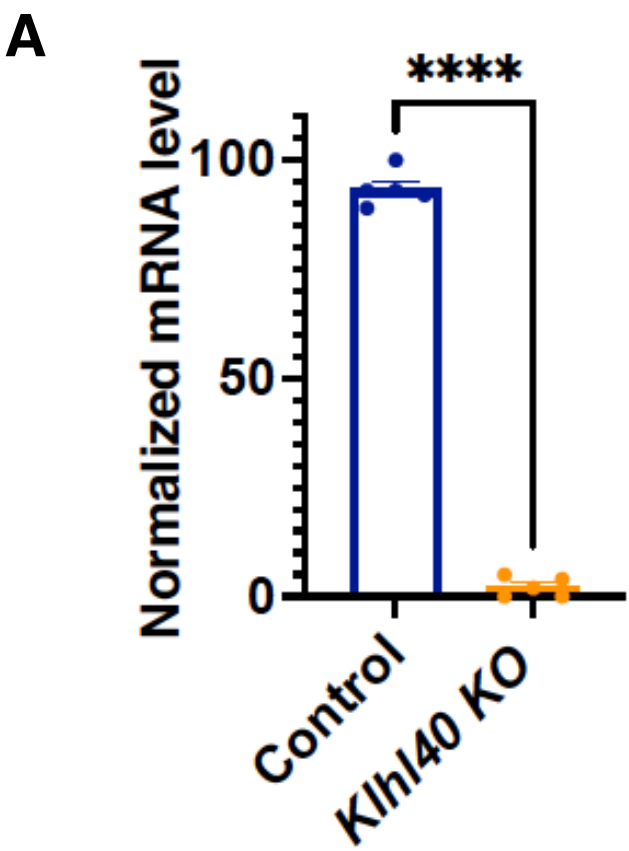

**B**

| Litter (#) | Wild-type (+/+) | Heterozygous (+/-) | Mutant (-/-) |
| --- | --- | --- | --- |
| 1 | 33.3% | 44.4% | 22.2% |
| 2 | 44.4% | 33.3% | 22.2% |
| 3 | 44.4% | 33.3% | 22.2% |
| 4 | 21.4% | 57.1% | 21.4% |
| 5 | 27.3% | 45.5% | 27.3% |

### Figure S2

**Figure S2**

**<u>Control</u> <u>*Klhl40* KO</u>**

**WB: KLHL40**

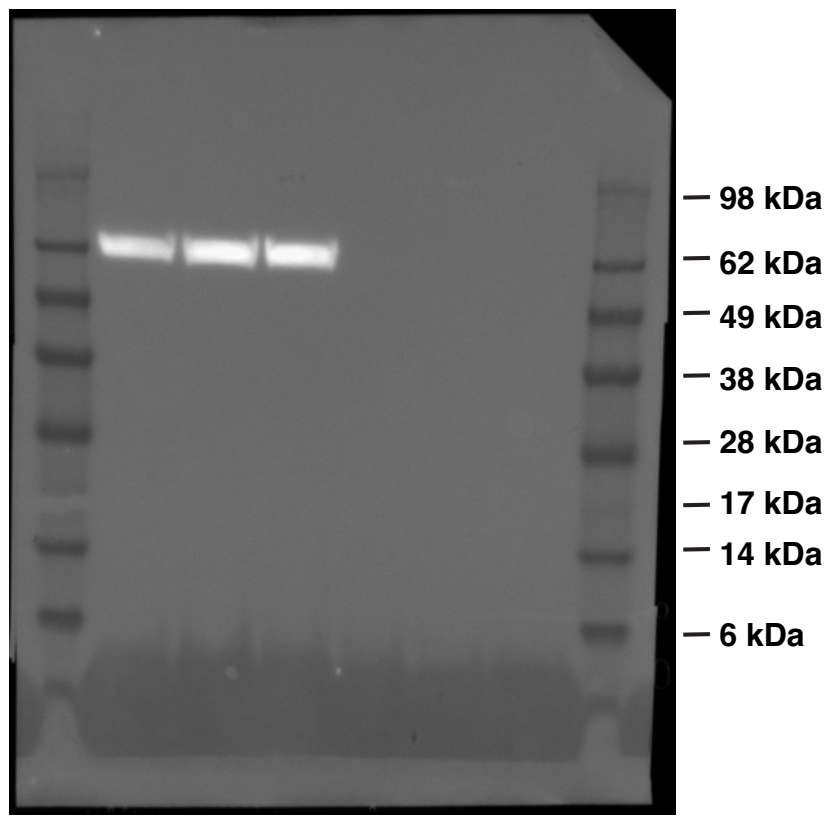

**<u>Control</u> <u>*Klhl40* KO</u>**

**WB: GAPDH**

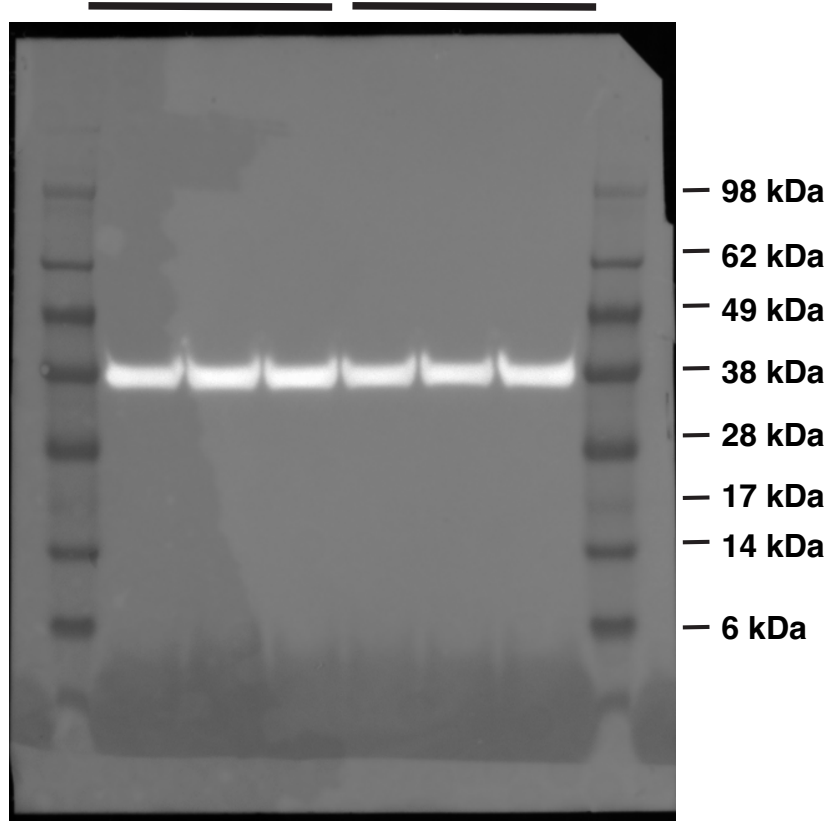

### Figure S3

**Figure S3**

**WT Control**

***Klhl40* KO**

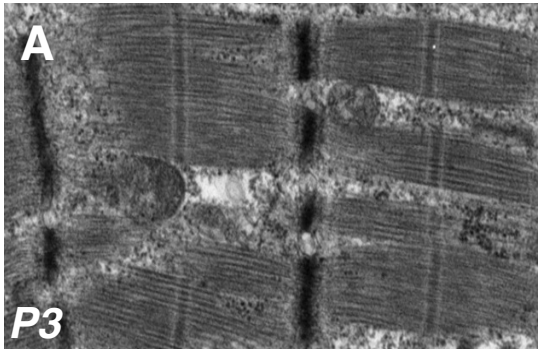

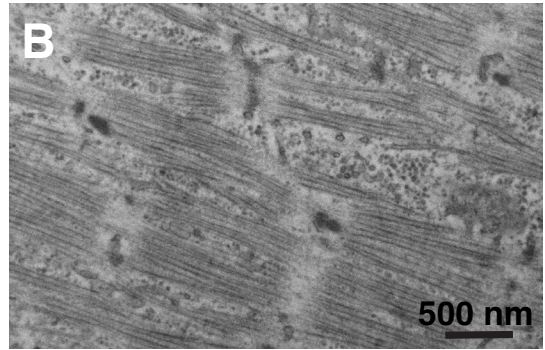

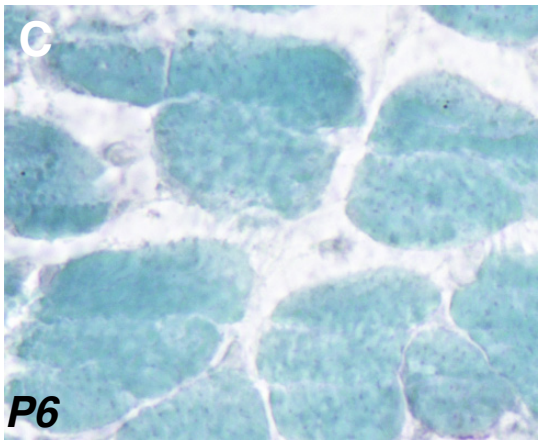

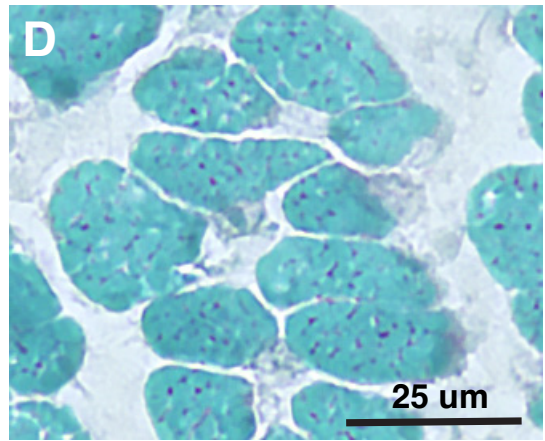

### Figure S4

**Figure S4**

**WT Control**

**A**

**WB: KLHL40**

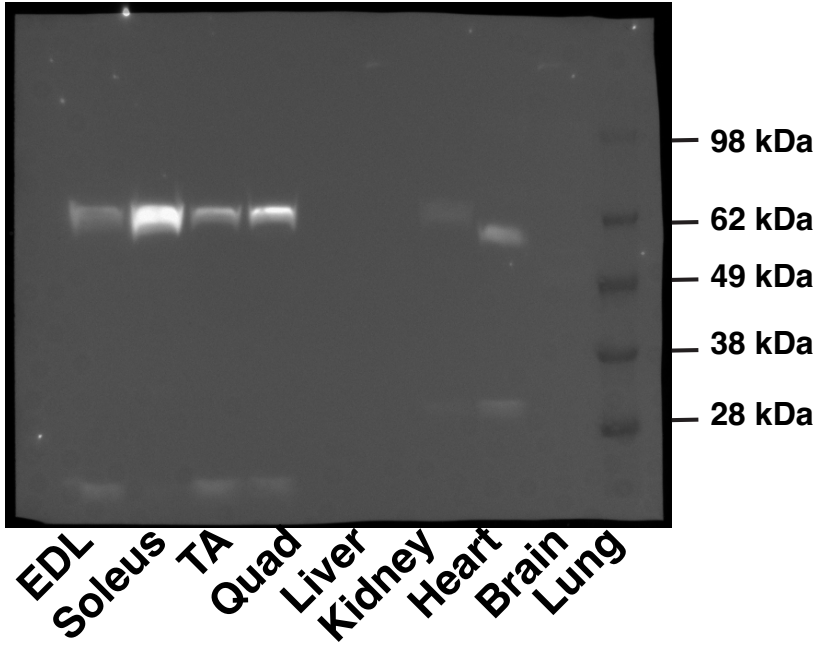

**WB: GAPDH**

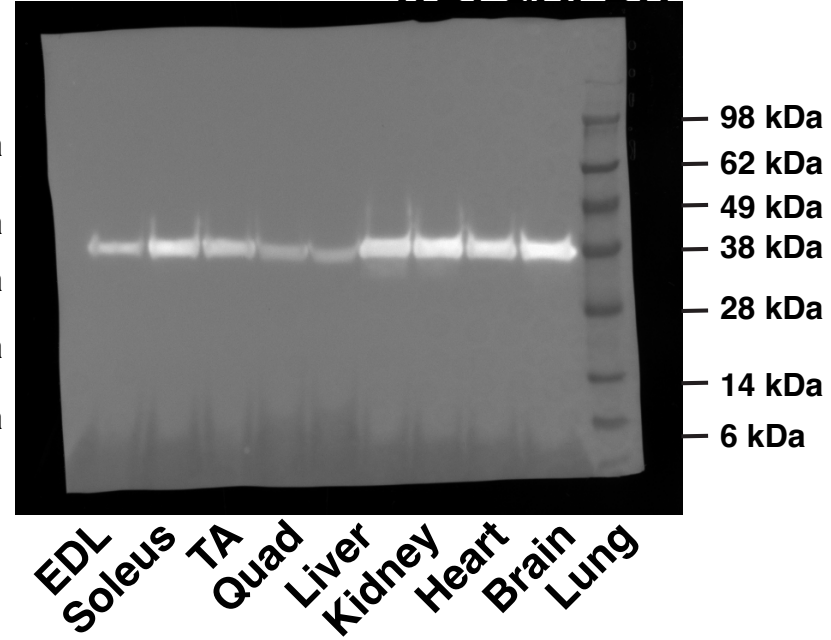

***Klhl40* KO treated**

**B**

**WB: KLHL40**

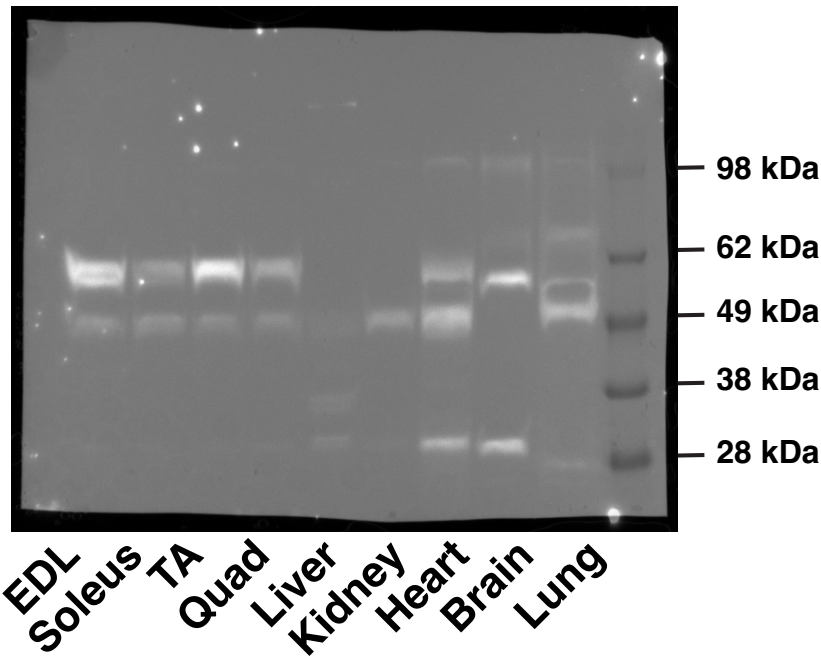

**WB: GAPDH**

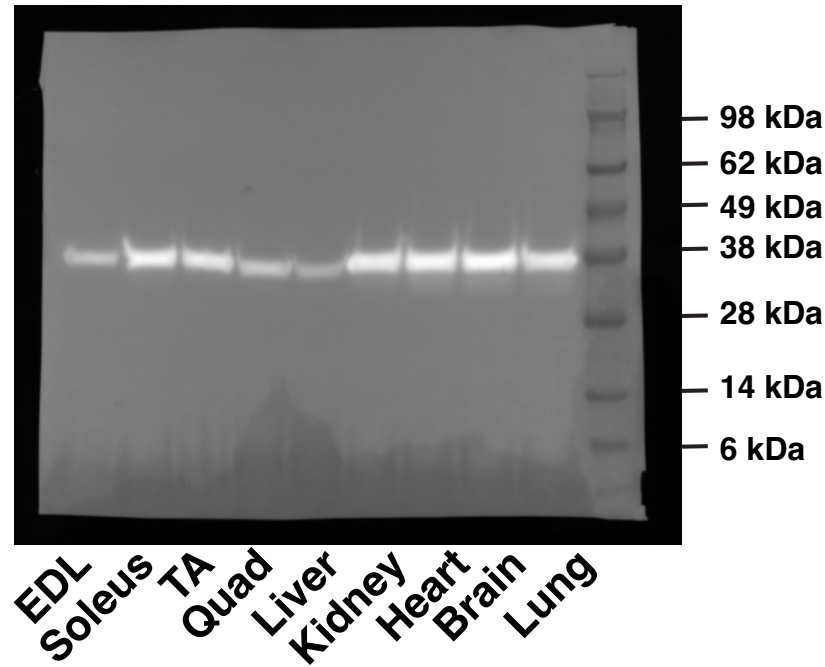
